# The protein folding rate and the geometry and topology of the native state

**DOI:** 10.1101/2021.10.06.463425

**Authors:** Jason Wang, Eleni Panagiotou

## Abstract

Proteins fold in 3-dimensional conformations which are important for their function. Characterizing the global conformation of proteins rigorously and separating secondary structure effects from topological effects is a challenge. New developments in Applied Knot Theory allow to characterize the topological characteristics of proteins (knotted or not). By analyzing a small set of two-state and multi-state proteins with no knots or slipknots, our results show that 95.4% of the analyzed proteins have non-trivial topological characteristics, as reflected by the second Vassiliev measure, and that the logarithm of the experimental protein folding rate depends on both the local geometry and the topology of the protein’s native state.

## Introduction

Proteins attain a specific conformation in space, called the native state, in order to perform their biological function. The process by which a protein attains its native state is called protein folding. Even though it is very difficult to experimentally see this process, it is possible to measure how fast a protein folds, the protein folding rate. Protein folding rates span many orders of magnitude. The topomer search model suggests that proteins “search” for their native state through an ensemble of possible conformations and that folding rate is determined by the ability of the protein to reach the topology of its native state^1^. This model emphasizes the importance of the 3-dimensional structure (also known as the tertiary structure) of the native state. It is natural therefore to hypothesize that the more complex a native state is, the slower its conformation will be attained, and thus, slower the folding rate of the protein. In this manuscript, we use rigorous tools from topology to characterize the complexity of the native state (in the absence of knots), and examine the role of topology and geometry on protein folding rates.

Many measures have been used to characterize the tertiary structure of the native state and its effect in protein folding^2–21^. One of the simplest characterizations of the native state is the number of sequence distant contacts, which is the number of sequence distant amino acids which are close in 3-space^2^. This quantity has shown one of the best correlations with experimental folding rates, suggesting that it captures something relevant to protein folding. However, it has been difficult to create a model of protein folding based on the number of sequence distant contacts alone. The number of contacts may in fact be a proxy for a more meaningful characteristic of a 3-dimensional conformation of a protein^8^.

A rigorous framework to define conformational complexity of curves is given in Knot Theory, which focuses on studying simple closed curves in 3-space (knots). Topological invariants are functions defined on closed curves which can classify them in different knot types^22^. Most efforts that aim at applying rigorous notions of topology to proteins, focus on identifying knots in proteins^23–31^. Proteins, however are not closed curves; by ignoring the chemical details and simply representing a protein by its CA atoms, the native state of a protein can be seen as an open ended polygonal curve in 3-space. Previous efforts to define knotting in proteins have relied on approximating the protein by a knot (or a knotoid, which is an open knot diagram). This method was very successful and revealed that many proteins contain knots or slipknots, but it also showed that knotted or slipknotted proteins comprise less than 10% of analyzed proteins^27,28,32–37^. Using this method, the rest of the proteins, are all assigned a trivial topological characterization. However, the native states of the remaining proteins are not identical and their folding rates, even when comparing single domain two-state proteins, differ over many orders of magnitude. It is therefore necessary to find new ways to characterize the tertiary structure of proteins that bridge the notion of topological complexity continuously from unknotted to knotted states. A measure of complexity which does not require an artificial closure of the proteins to measure protein complexity is the Gauss linking integral. When applied over a protein or an arc of a protein, it gives the Writhe or the Average Crossing Number. It can also be used to study the linking between parts of a protein. These measures have shown strong correlation with protein folding rate and support the hypothesis that a more geometrically/topologically complex native structure leads to a lower folding rate^3,38^. However, the Writhe and the ACN are significantly affected by the local geometry of the protein and are not strong measures of topological complexity.

To decouple geometry from topology of proteins and capture topological effects in both knotted and unknotted proteins, with or without slipknots, we propose a new measure of the protein complexity; the second Vassiliev measure^39^. We show that the second Vassiliev measure can be applied to proteins without artificial closure of the chains to quantify topological complexity (including knotting). It takes a non-trivial value for most proteins and reflects various subtle degrees of topological complexity, varying continuously from trivial to knotted topology. In contrast to the Writhe and ACN, this tool is not affected by secondary structure elements. We apply the Writhe, the Average Crossing Number (ACN) and the second Vassiliev measure to a set of two-state proteins (which are known to fold in an all or none fashion) and a set of multi-state proteins (known to have well populated intermediates). Our results show that these measures capture different characteristics of the tertiary structure of the native state and that the folding rate depends both on the geometry and the topology of the native state, even for proteins without knots or slipknots.

## 1 Results

In this Section, we analyze the geometry and topology of a data set of two-state and multi-state proteins whose folding rates were reported in^16^. Each protein was represented as a polygonal curve by connecting the consecutive alpha carbon atoms, CA atoms, with a line segment. The coordinates of the CA atoms were obtained from the Protein Data Bank (PDB)^40^.

The Writhe, we denote *Wr*, and the Average Crossing Number, *ACN*, are derived from the Gauss linking integral and are very sensitive on the local geometry of proteins. The second Vassiliev measure, we denote *v*_2_, and the second Absolute Vassiliev measure, we denote *Av*_2_, are topological measures, quantifying the overall 3-dimensional structure, and are not as sensitive on the local geometry. Higher values of Writhe and ACN in random polygonal curves are in principle associated with higher topological complexity. In proteins however, high values of Writhe or ACN may not necessarily reflect topological complexity of the backbone, as they are significantly affected by the presence of secondary structure elements. In particular, helices contribute high values of Writhe and ACN to the total Writhe and ACN of the protein. Previous work has shown that experimental folding rates correlate with both the Writhe and the ACN of the native state^8^, but it is difficult to decouple the role of topology from the role of geometry or the role of secondary structure elements.

To detect the effect of topology in protein folding, we propose using a stronger measure of topological complexity, the second Vassiliev measure, *v*_2_, and the absolute second Vassiliev measure, *Av*_2_. The latter is used to capture contributions to *v*_2_ that cancel out because of opposite signs. *v*_2_ (and *Av*_2_) is not as sensitive to the local geometry as the Writhe or ACN, allowing it to capture the characteristics of the global conformation of a protein (see Figure 1 for an illustrative example). In general, higher values of *v*_2_ represent higher knotting complexity. Proteins with no knots may have values of *v*_2_ that are much less than 1 in magnitude, but non-zero. Therefore, even though most proteins do not contain knots, they may be topologically not trivial, as it is reflected by non-zero values of *v*_2_.

**Figure 1.**
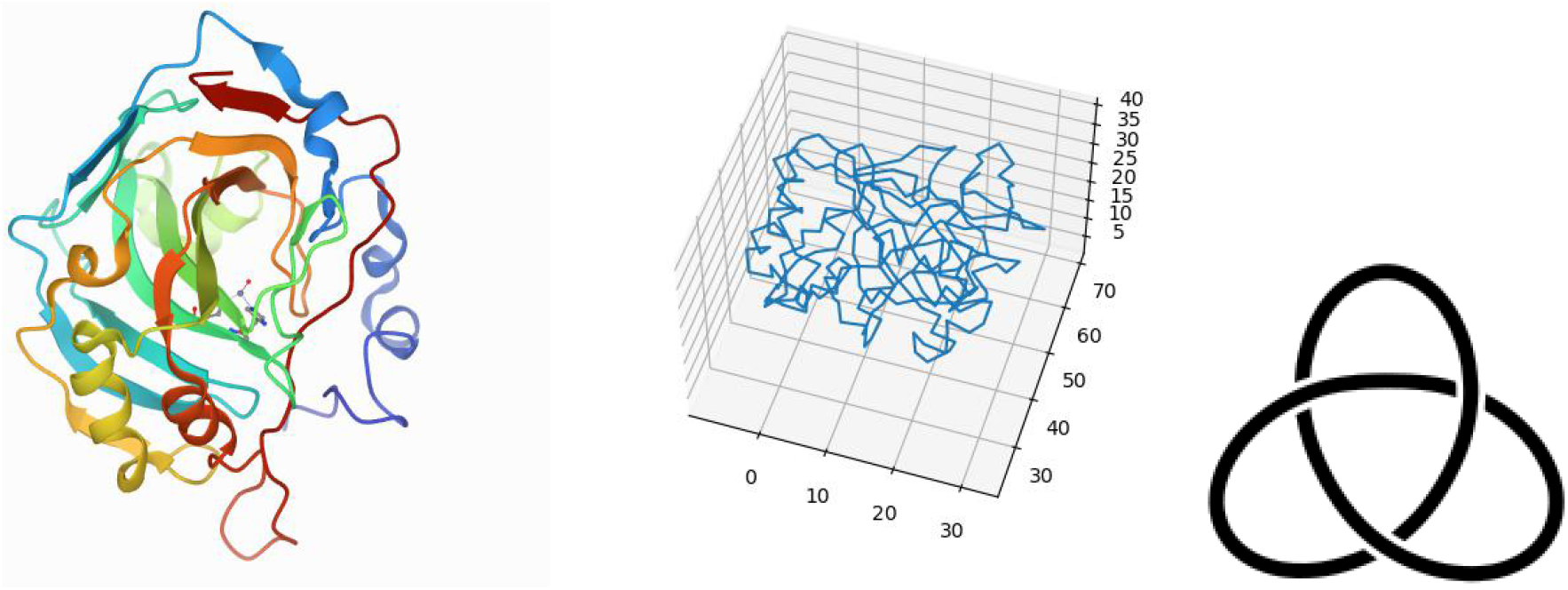
Protein 1v9e from the PDB (Left) and as a simple polygonal curve (Middle) and a mathematical trefoil knot (Right). 1v9e has *v*_2_ = 0.808 and *Av*_2_ = 0.844. Note that *v*_2_ = 1 corresponds to the trefoil knot. Indeed, 1v9e is known to contain a trefoil knot using the knot fingerprint approach^27^. Its *ACN* = 117.6653 and *Wr* = 6.28669.

Both *v*_2_ and *Av*_2_ quantify how “knotted” the tertiary structure of a protein is or has the potential to be, while *Wr* and *ACN* are measures of entanglement complexity of the protein, including secondary structure element effects. These can have different values on the same protein. Examples of unknotted proteins with similar ACN and Writhe values but different *v*_2_ values are shown in Figure 2.

**Figure 2.**
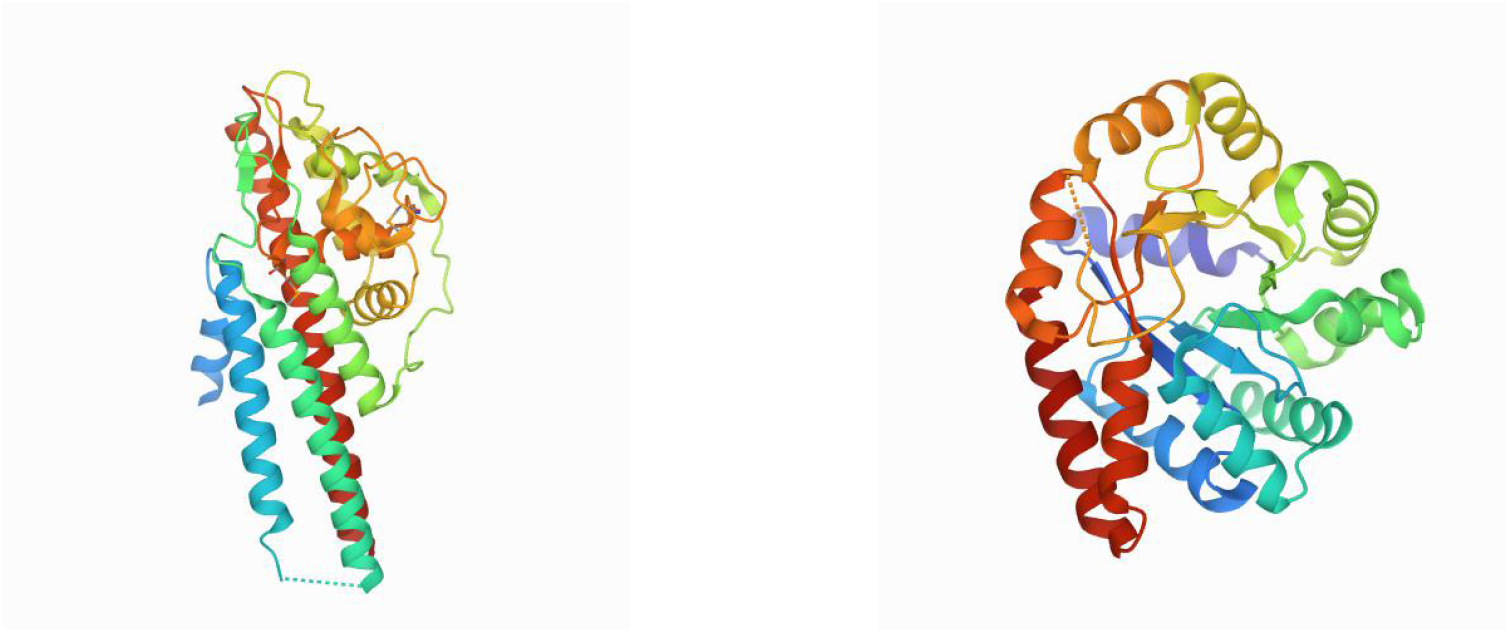
Protein 1l8w (Left) and protein 1qop (Right). They both have similar Writhe and ACN but different *v*_2_ values. Namely, Protein 1l8w has *Wr* = 28.27, *ACN* = 143.61 and *v*_2_ = 0.022. Protein 1qop has *ACN* = 131.5524,*Wr* = 23.00797 and *v*_2_ = 0.229.

In Section 1.1 we present our results on the topology and geometry of a small set of proteins. In Section 1.2 we present our results on folding kinetics and topology of the native state for a mixed set of two and multi-state proteins. In Section 1.3 we focus only on the two-state proteins and in Section 1.4 we focus only on the multi-state proteins.

### 1.1 The topology and geometry of proteins

We analyze a set of proteins with no knots and no slipknots. Such proteins have been “invisible” by the common mathematical methods to characterize protein conformations. However, we find that 95.4% of the proteins analyzed in this study have non-zero value for the second Vassiliev measure. Putting this result in the context of previous studies, we note that by using the knot fingerprint method and the HOMFLYPT polynomial, it has been shown that less than 10% of proteins in the PDB contain a knot^28^, while other studies show that 32% of proteins contain an “entangled motif” (two disjoint subchains with a Gauss linking integral of magnitude greater than 1)^41^. Here we provide a simple measure of topological complexity that applies to practically all proteins to characterize their topological complexity.

We apply the second Vassiliev measure, as well as the Writhe and ACN on a set of multi-state and two-state proteins. We stress that these measures capture different conformational information. Figure 3 shows the correlation of the different topological measures used in this study. We see that the Writhe and ACN capture different information than the *v*_2_ and *Av*_2_. Indeed, even though the Writhe and the ACN are also measures of conformational complexity, related to topology, they are more sensitive to local entanglement rather than topology. In particular, Writhe and ACN are impacted by secondary structure elements.

**Figure 3.**
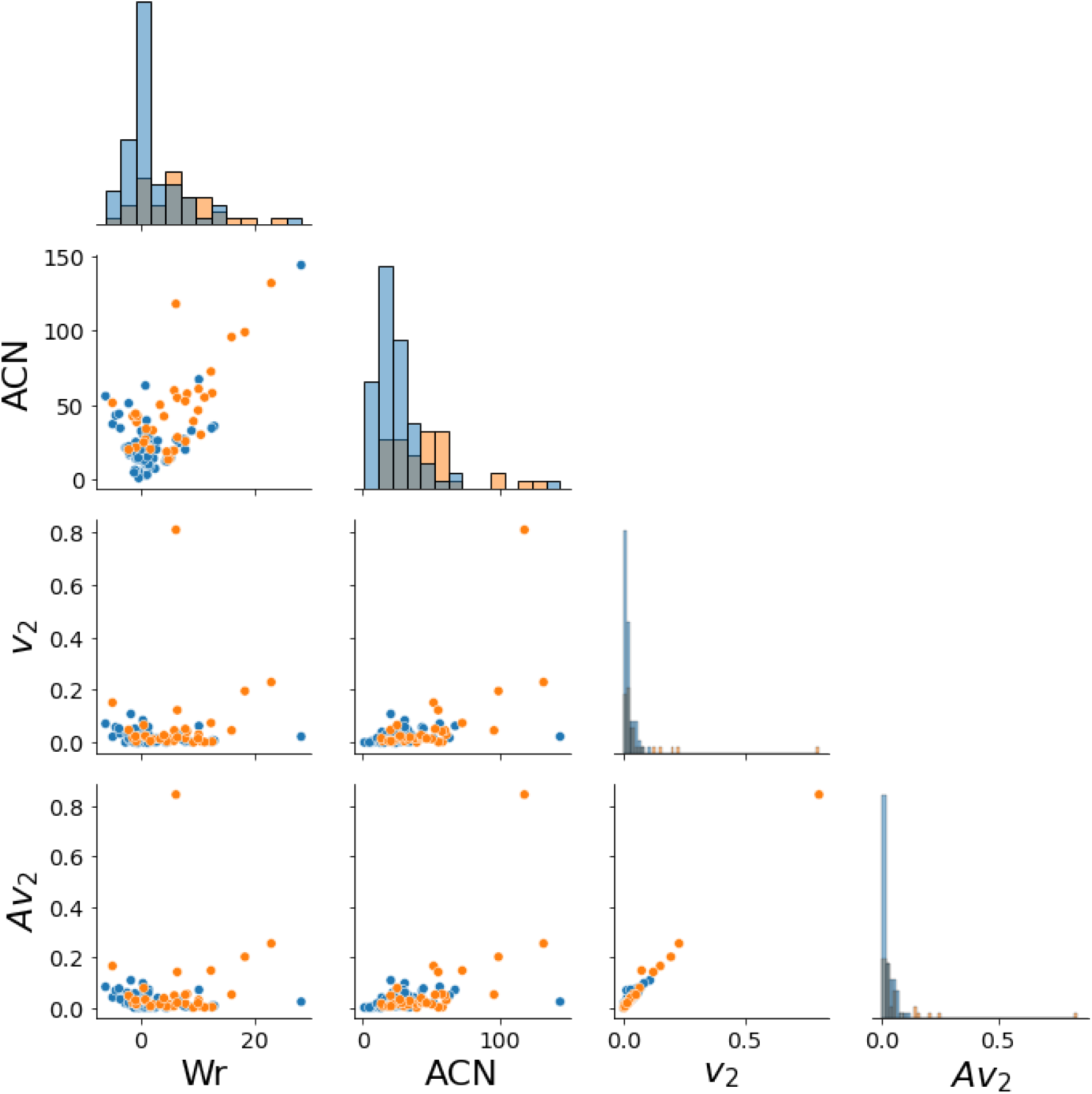
The plots on the diagonal show the distribution of each measure for the protein set. The scatter plots compare the measures pairwise for each protein. Blue points represent two-state and orange points represent multi-state proteins. We see that there is little correlation between Wr and *v*_2_, Wr and *Av*_2_, ACN and *v*_2_, or ACN and *Av*_2_. Wr and ACN show a correlation, as well as *v*_2_ and *Av*_2_.

### 1.2 Folding rate as a function of the geometry and topology of the native state for a mixed (two-state and multi-state) set of proteins

Figure 4 shows the logarithm of the experimental folding rate as a function of the Writhe (Left) and normalized Writhe (Right) of the native state. We see a correlation *r* = 0.119 and *r* = 0.364, respectively, and the slope of the regression line is positive, which seems to contradict the hypothesis that more complex folded structures would be achieved at a slower rate. Similar results were observed for a set of two-state proteins in^8^. At a closer inspection, we see that the folding rate decreases with more negative Writhe values. This was also observed in^8^. This further corroborates the result that the Writhe captures some aspects of handedness related to folding rates, as well as secondary structure elements. In particular, we know that helices contribute a positive value of Writhe. In an effort to decouple the local secondary structure effect from the topology of the protein using the Writhe, the Writhe of the primitive path was introduced in^8^ and it was shown that the folding rate correlates better with the latter. Note that previous results have also showed a different impact on folding rate of local versus global properties of the protein^42^.

**Figure 4.**
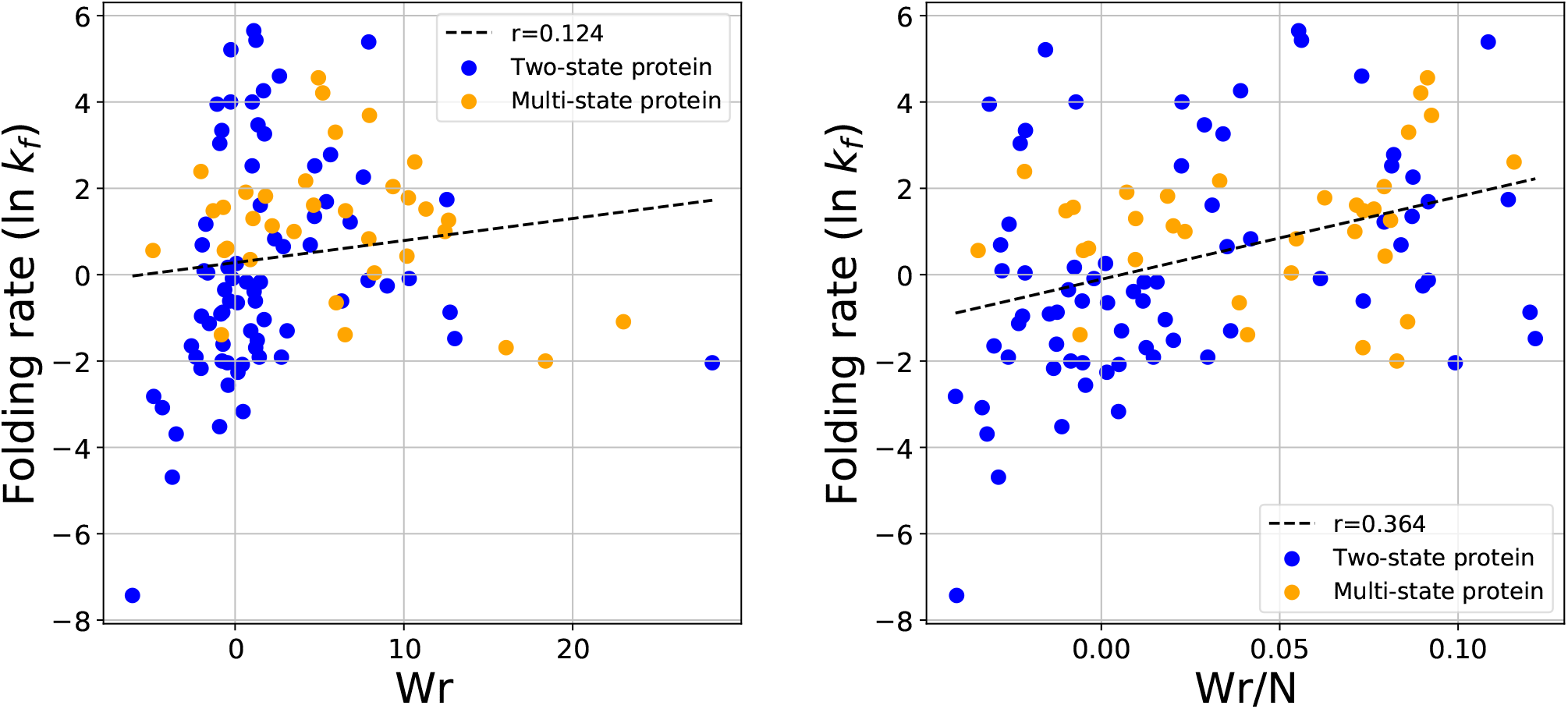
The protein folding rate as a function of the writhe (left) and writhe/N (right) for all proteins in the data set. The folding rate is represented as the natural log of the experimental folding rate *k*_*f*_.

Figure 5 shows the logarithm of the experimental folding rate as a function of the ACN (Left) and normalized ACN (Right). We notice that the folding rate decreases with increasing ACN and ACN/N with a correlation, *r* = −0.4 and *r* =− 0.5, respectively. This agrees with the hypothesis that proteins fold slower to more complex native states. However, the fact that the folding rate decreases with ACN it does not mean necessarily that it is affected by the topological complexity of the tertiary structure of the protein. The ACN, like the Writhe, is affected by the presence of secondary structures. So, the question remains to what extent the folding rate depends on the global topology versus the local entanglement of the native state.

**Figure 5.**
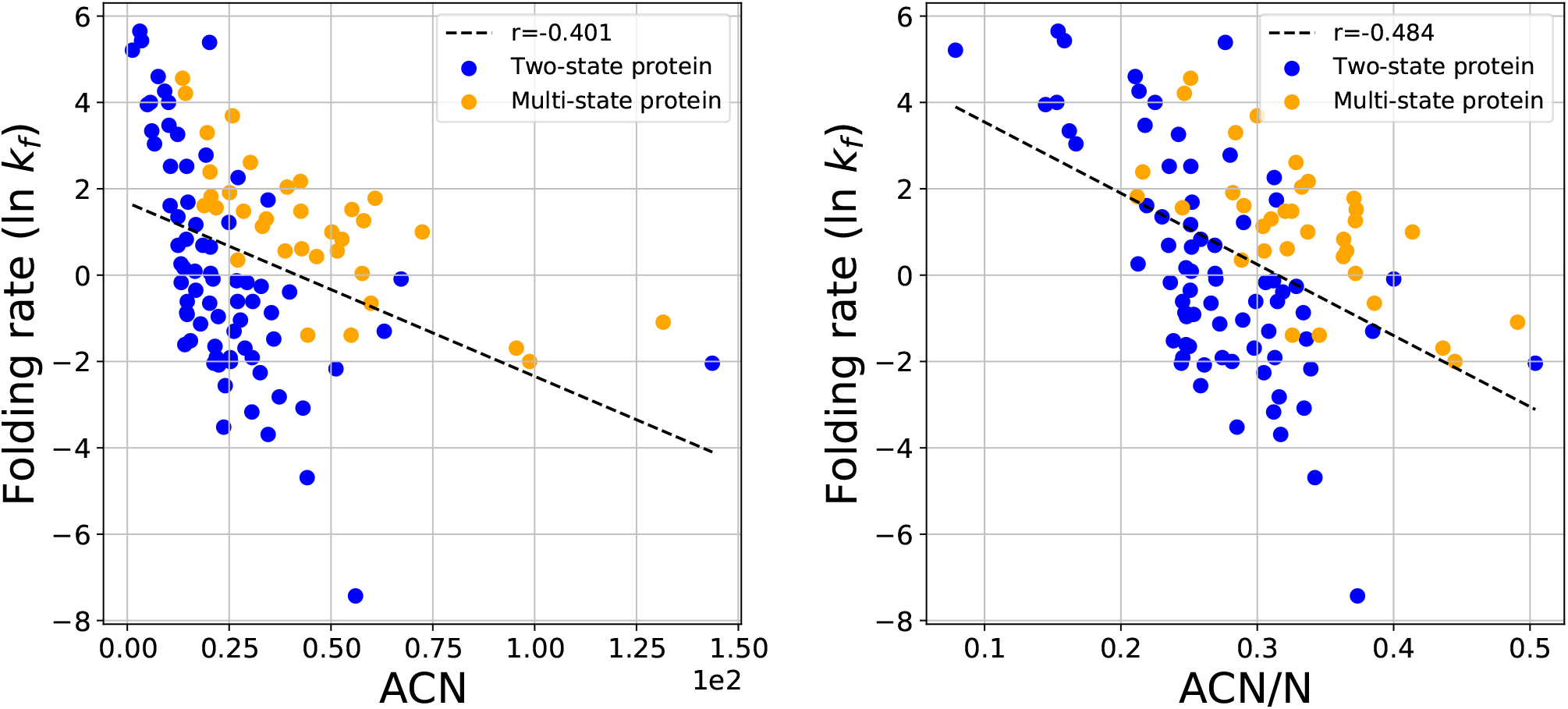
The folding rate as a function of the native state ACN (left) and the ACN/N (right). Multi-state proteins tend to have higher ACN than two-state proteins with similar folding rates.

The second Vassiliev Measure (*v*_2_) is a better indicator of topology, measuring topological complexity in the global conformation rather than being affected by local entanglement. Figure 6 shows the logarithm of the experimental folding rate as a function of *v*_2_ (Right) and *v*_2_/*N* (Left). We see that the logarithm of the experimental folding rate decreases with increasing *v*_2_ with *r* = −0.361 and −0.364, respectively. This result shows that the folding rate decreases with the topological complexity of the native state. Note that this is the first result that shows that folding rate correlates with aspects of global complexity, and specifically topology, irrespective of local structure.

**Figure 6.**
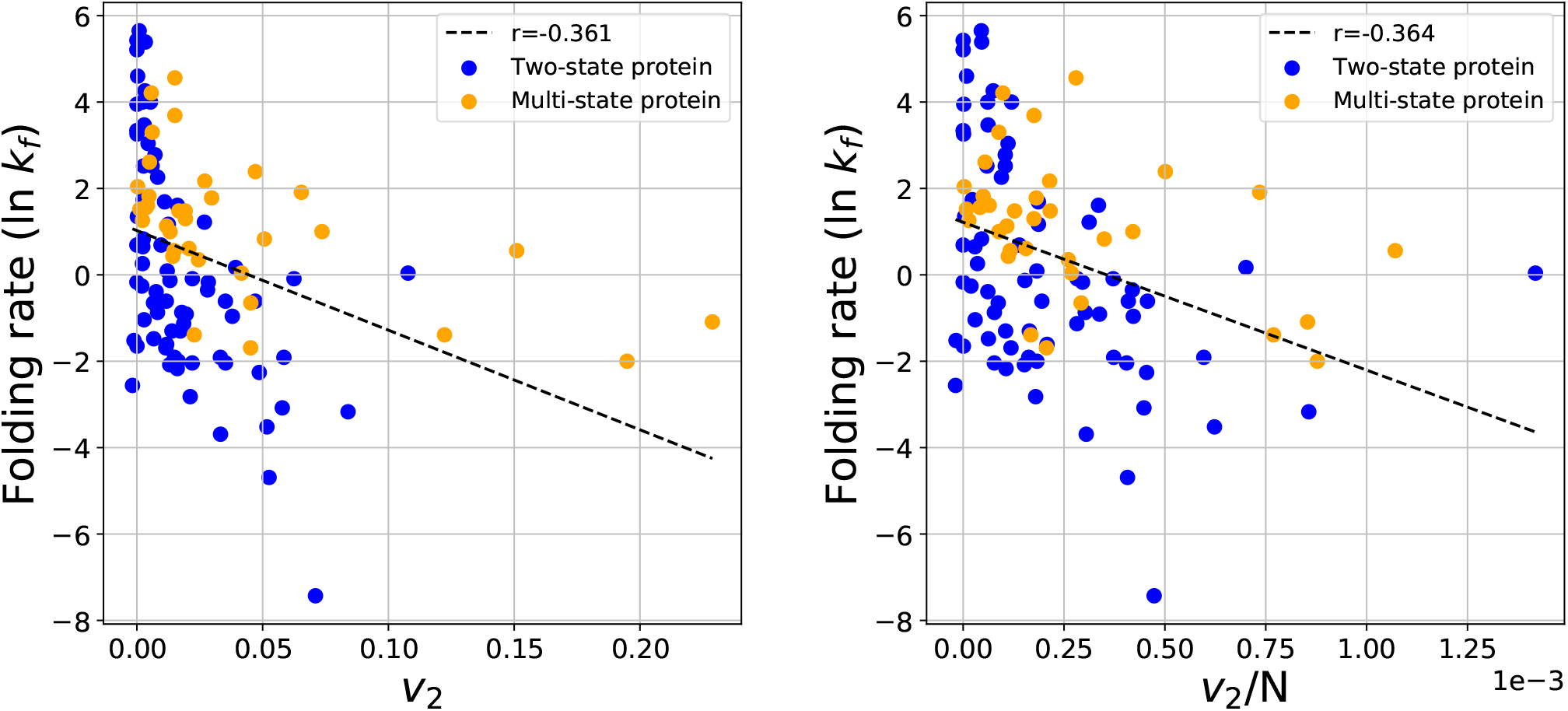
The protein folding rate as a function of *v*_2_ (left) and *v*_2_/N (right) for all proteins.

Figure 7 shows the logarithm of the experimental folding rate as a function of *Av*_2_. Our results show that the folding rate has a correlation of order *r* = −0.398 and *r* = −0.431 with *Av*_2_ and *Av*_2_/*N*, respectively. This is in agreement with the results on *v*_2_.

**Figure 7.**
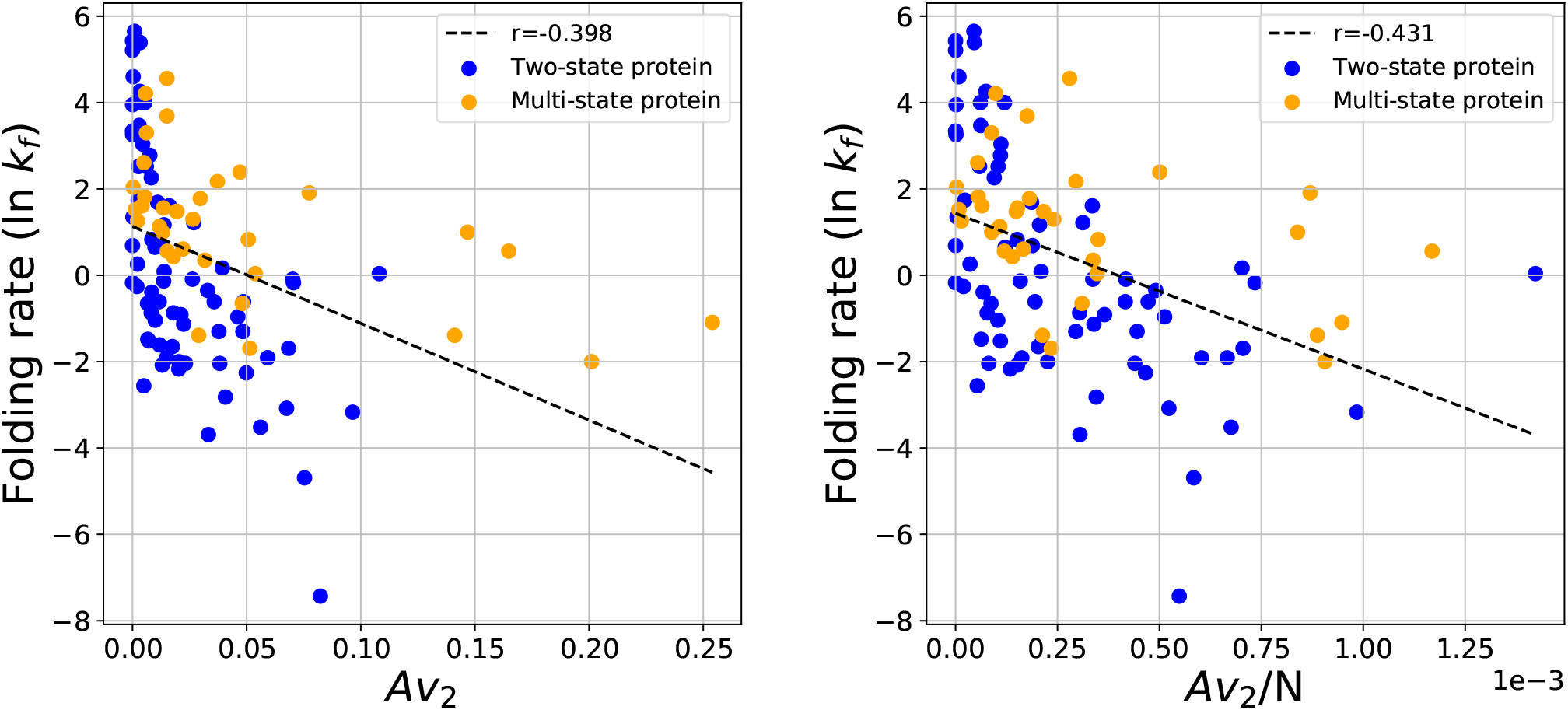
The protein folding rate as a function of the approximate *Av*_2_ (left) and *Av*_2_/N (right) for both two-state and multi-state proteins.

### 1.3 Two-State Proteins

In the set of two-state proteins, as in the combined set, the logarithm of the experimental folding rate shows the strongest correlation with ACN and ACN/N, with coefficients of *r* = −0.522 and *r* = −0.653, respectively, as seen in Figure 8.

**Figure 8.**
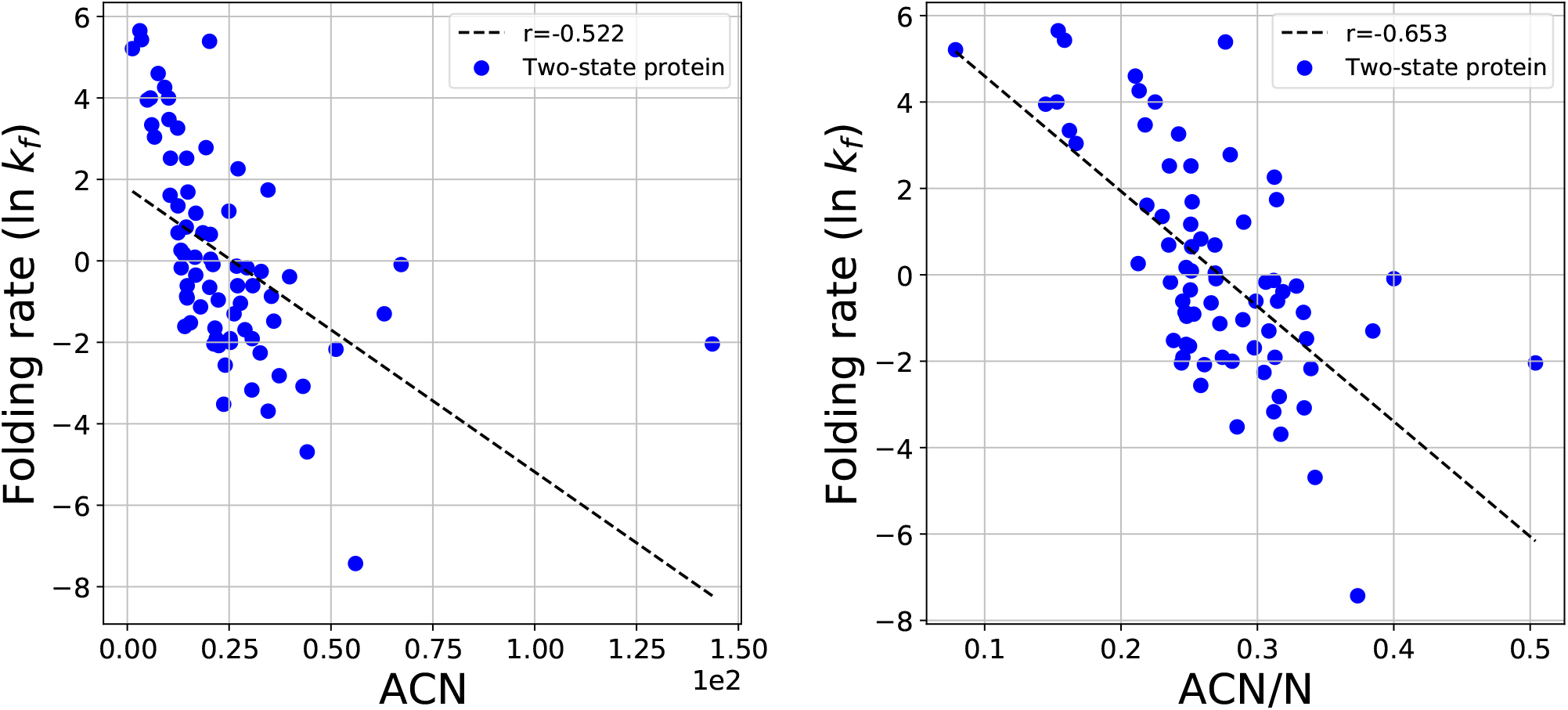
The protein folding rate as a function of the ACN (left) and ACN/N (right) of the set of two-state proteins. This set excludes the protein 1l8w which had an outlying ACN of about 140. When included the correlation decreases to r = −0.48.

The experimental folding rate of two-state proteins and the Writhe and Writhe/N shows a correlation *r* = 0.309 and *r* = 0.369, respectively and it is shown in the SI.

We find that the folding rate shows a stronger correlation with *v*_2_ of the two-state proteins alone, compared to the mixed set. Figure 9 shows the logarithm of the experimental folding rate as a function of *v*_2_ and *Av*_2_ for two-state proteins. We find that the logarithm of the experimental folding rate decreases with increasing *v*_2_ and *Av*_2_ values with *r* = −0.546 and *r* = −0.611, respectively.

**Figure 9.**
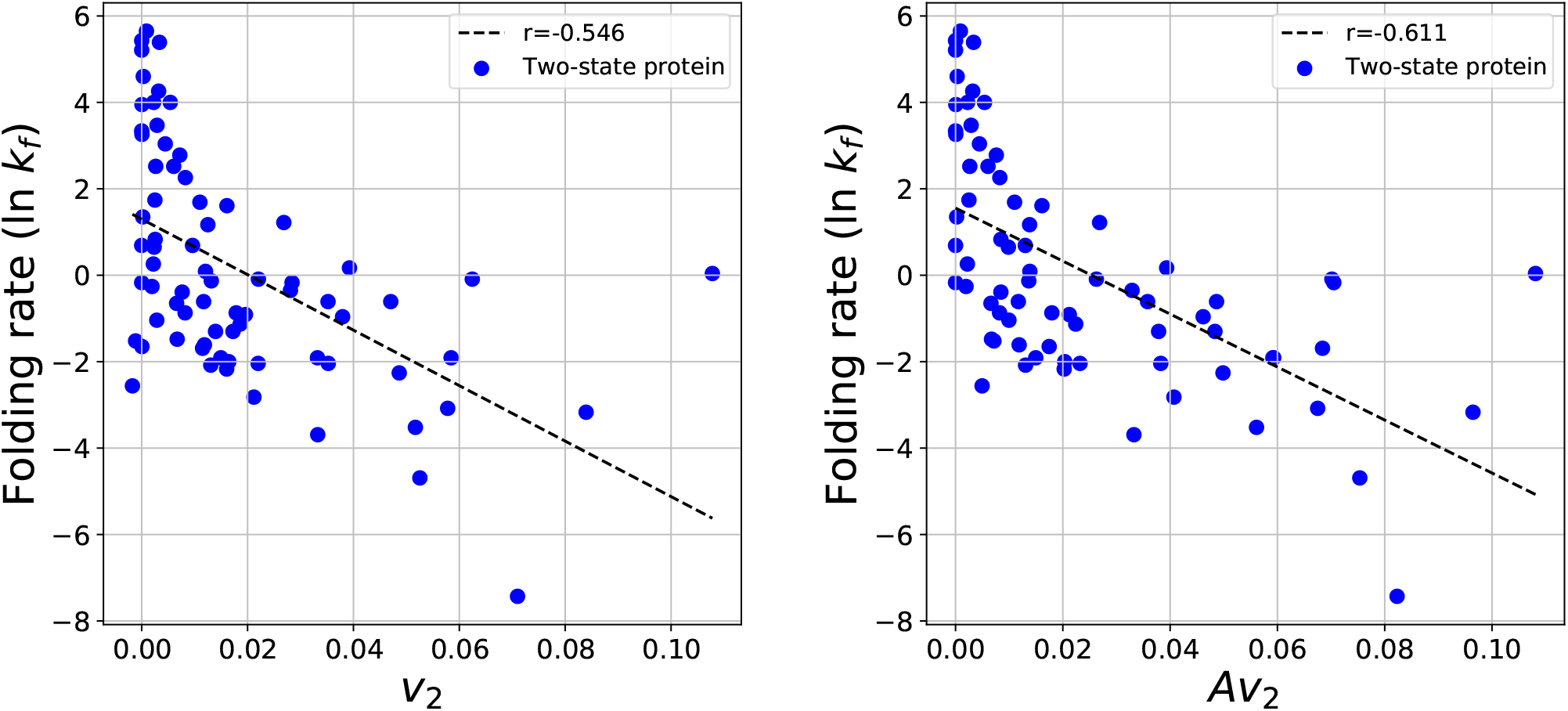
The protein folding rate as a function of the *v*_2_ (left) and *Av*_2_/N (right) of the set of two-state proteins.

### 1.4 Multi-state Proteins

In the set of multi-state proteins, the logarithm of the experimental folding rate shows a particularly strong correlation with ACN, with *r* = −0.727 and *r* == 0.668, shown in Figure 10.

**Figure 10.**
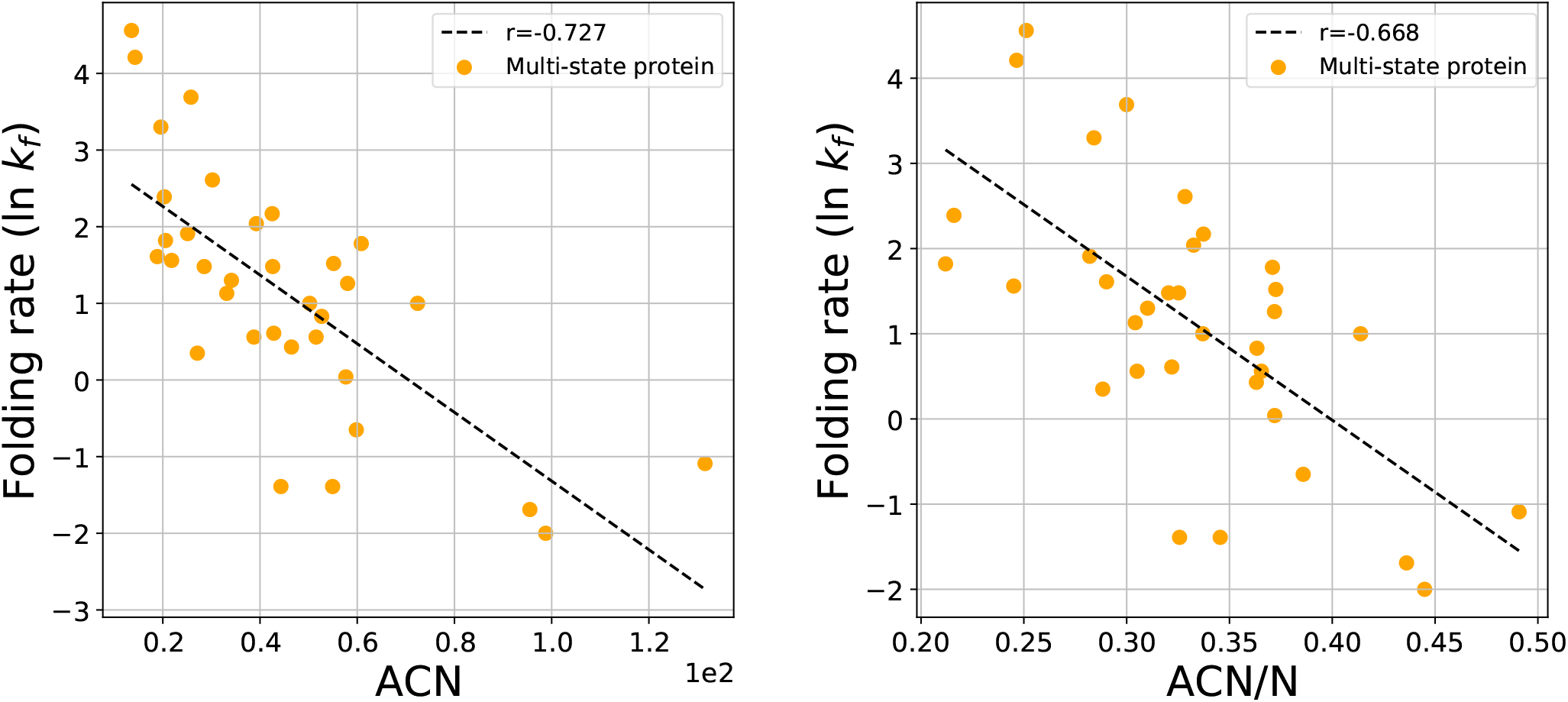
The protein folding rate as a function of the ACN and ACN/N of the set of multi-state proteins.

The logarithm of the experimental folding rate shows a weak correlation with the Writhe and Writhe/N; *r* = 0.182 and *r* = 0.220 (data shown in the SI).

Figure 11 depicts the protein folding rate against *v*_2_ and *Av*_2_ for multi-state proteins. The logarithm of the experimental folding rate shows a strong correlation with these measures with *r* = −0.563 for *v*_2_ and *r* = −0.554 for *Av*_2_.

**Figure 11.**
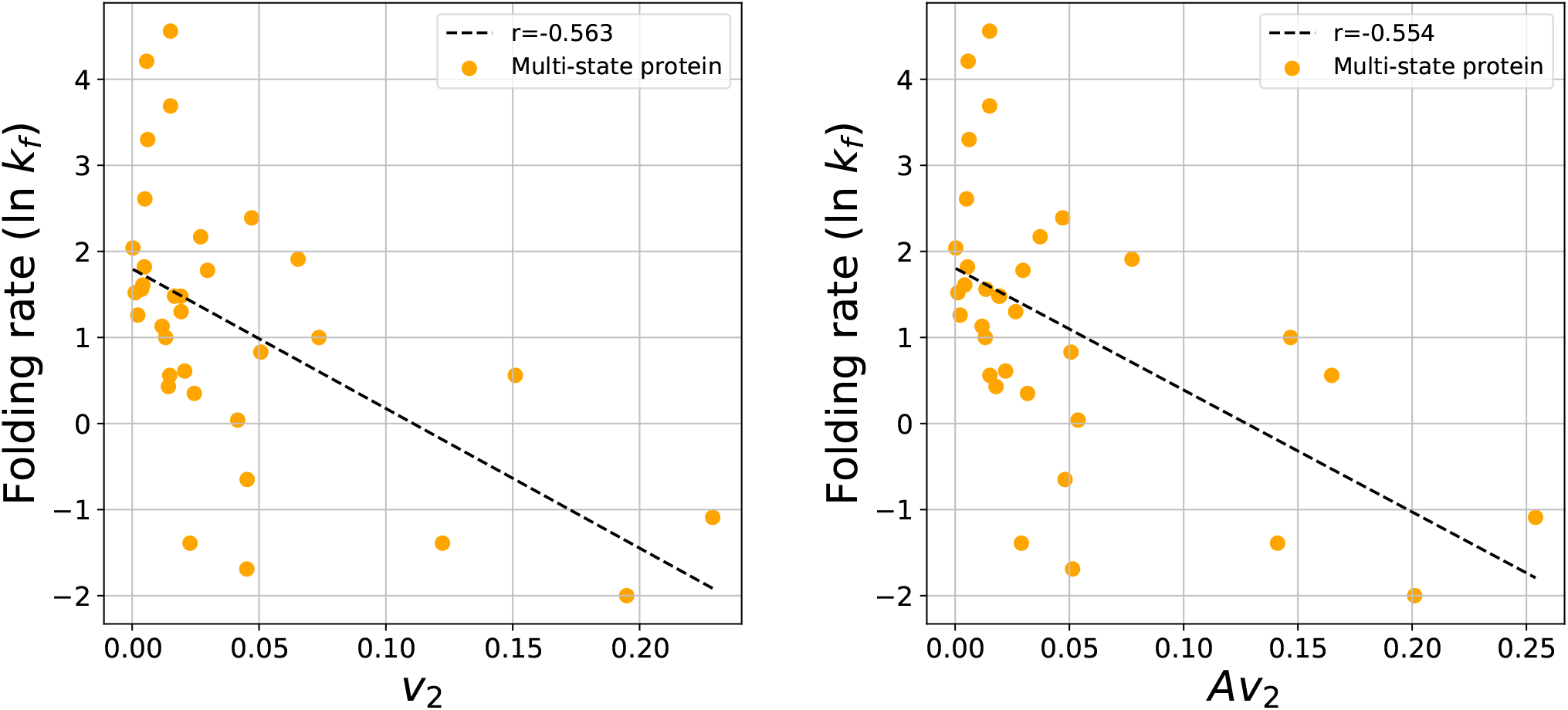
The protein folding rate as a function of the *v*_2_ and *Av*_2_ of the set of multi-state proteins.

## Discussion

Many previous measures have been proposed to quantify the complexity of the native state of proteins. Even though the folding rate correlates strongly with some simple measures, none of them proves that the topology of the 3-dimensional conformation of the entire protein plays a role in protein folding. In this manuscript, we show that indeed folding rates depend on the mathematical topology of the native state, even for unknotted proteins. This is done using the second Vassiliev measure, a new measure of topological complexity of proteins that can characterize the topology of protein conformations continuously from trivial to knotted state.

Our results on the correlation of the logarithm of the experimental folding rate and the Writhe, ACN and the second Vassiliev measure, *v*_2_ and *Av*_2_, of the protein native state show that the folding rate depends on both the geometry of the native state, as well as its topology.

We also find that the folding rate correlates better with the normalized *ACN*, indicating a strong correlation between *ACN* and the size of the protein, a parameter already known to impact the protein folding rate^12,43−45^. On the other hand, *v*_2_ does not show any improvement upon normalizing by the length of the protein. This also corroborates the fact that *v*_2_ captures topological aspects of the tertiary structure of the protein, which are independent of its length.

Looking at the sets of two-state proteins or multi-state proteins individually, each measure has a stronger correlation with the folding rate than in the combined set. This may be expected, as two-state and multi-state proteins have different folding mechanisms^46^. It is interesting to point out that the folding rate of two-state proteins shows a stronger correlation with *v*_2_, which is a purely topological measure, than with *ACN*, which is affected strongly by local geometry. The folding rate of multi-state proteins, however, shows a stronger correlation with *ACN* than with *v*_2_. This may suggest that local structure is more involved in the folding mechanism of multi-state proteins, compared to that of two-state proteins.

It is possible that many of the proposed measures together could provide better correlations with protein folding rate^9^. However, in addition to correlation, it is important to establish causation. Our results show that the topology of the native state, as it is captured by rigorous and well understood mathematical tools from knot theory, should be accounted in a model of protein folding.

## Methods

In this Section, we give the definitions of the mathematical measures used to characterize the 3-dimensional conformation of proteins.

The Writhe of a curve in 3-space is defined as the Gauss linking integral over the curve^47^:

### Definition 1.1

*For an oriented curve 𝓁 with arc-length parametrization γ*(*t*), *the Writhe, Wr, is the double integral over l:*

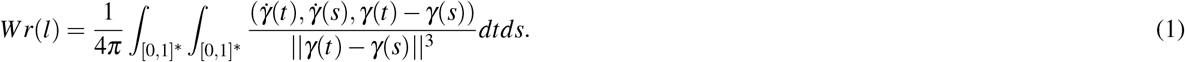

where integration is over all *s,t* ∈[0, 1], *s* ≠*t*.

The Writhe measures how much the chain turns around itself. Taking into account the orientation of the curve (from start to end-point), given a projection of the curve, one can add up the number of crossings, with signs according to orientation and the convention shown in Figure 12. The Writhe is a real number, equal to the average algebraic sum of crossings over all possible projection directions.

**Figure 12.**
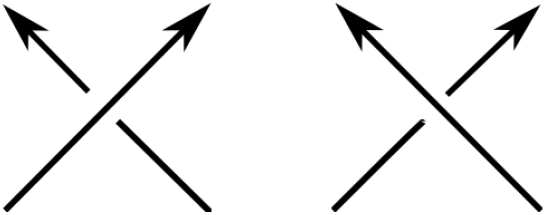
A crossing in a projection of a protein can be assigned a positive (Left) or negative (Right) sign, depending on the orientation.

By taking the absolute value of the integrand, we obtain the Average Crossing Number. Namely,

### Definition 1.2

*For an oriented curve 𝓁 with arc-length parametrization γ*(*t*), *the Average Crossing Number, ACN, is the double integral over l:*

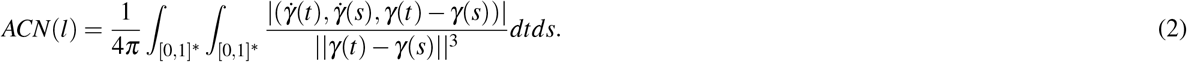

where integration is over all *s,t* ∈[0, 1], *s ≠ t*.

The ACN is a positive real number that measures the average sum of crossings (without signs) over all possible projection directions.

Both Writhe and ACN can be computed exactly, avoiding numerical integration, using the algorithm described in^48^. The Writhe and the ACN are continuous functions of the chain coordinates (not topological invariants) for both closed and open curves.

The second Vassiliev measure of open curves in 3-space was introduced in^39^ and is defined as follows:

### Definition 1.3

*For an oriented curve l with parametrization γ*(*t*), *v*_2_ *is defined using the following integral:*

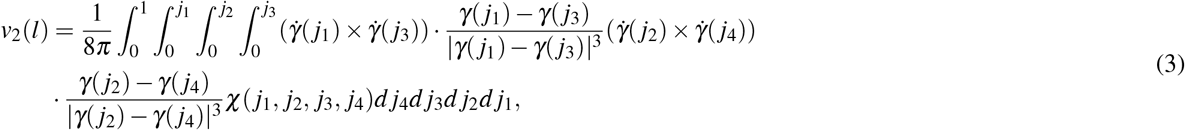

*where χ*(*j*_1_, *j*_2_, *j*_3_, *j*_4_) = 1, *when* (*j*_1_, *j*_2_, *j*_3_, *j*_4_) ∈ *E and χ*(*j*_1_, *j*_2_, *j*_3_, *j*_4_) = 0, *otherwise and where E* ⊂ [0, 1]^4^, *such that* Γ(*j*_1_, *j*_3_) = −Γ(*j*_2_, *j*_4_), *where* 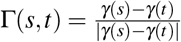, *for s,t* ∈ [0, 1].

The second Vassiliev measure of an open curve in 3-space is equal to the average of the algebraic sum of “alternating pairs” of crossings over all projection directions. Namely, for any given projection of the curve, an alternating pair of crossings, (*j*_1_, *j*_3_) and (*j*_2_, *j*_4_), is such that that if the projection of *γ*(*j*_1_) is over, resp. under, that of *γ*(*j*_3_), then the projection of *γ*(*j*_2_) is under, resp. over, that of *γ*(*j*_4_). The second Vassiliev measure does not have a closed form and can only be estimated as the average over a large number of projections. In this manuscript, *v*_2_ was estimated as an average over 10,000 projections for two-state and 5,000 projections for multi-state proteins.

For closed curves, the second Vassiliev measure is a second Vassiliev invariant of knots and it is an integer topological invariant that can distinguish several knot types. For proteins, the second Vassiliev measure is a real number that is a continuous function of the chain coordinates in 3-space, and, if the protein ties a knot, it tends to the topological invariant of the knot. We note that the term “topology” in mathematics may be elusive for open curves in 3-space. It would be more accurate to use another term, such as “potential topology” for such curves. However, in this manuscript, we will use the term topological complexity, for all proteins, when *v*_2_ ≠ 0.

Since *v*_2_ is an average algebraic sum of patterns of crossings in a projection over all projection directions, positive values in one projection may cancel with negative values in another. For this reason, we introduce *Av*_2_, which we define by taking the absolute value in the integrand in Eq. 3. For open curves, this is a positive number, which varies continuously with the coordinates of the chain and as the endpoints of the chain tend to coincide, it tends to the absolute second Vassiliev invariant of the resulting knot.

## Acknowledgements

JW and EP acknowledge support from NSF REU 1852042, NSF DMS 1913180, NSF 1925603 and NSF CAREER 2047587.

